# Feedback-based motor control can guide plasticity and drive rapid learning

**DOI:** 10.1101/2022.10.06.511108

**Authors:** Barbara Feulner, Matthew G. Perich, Lee E. Miller, Claudia Clopath, Juan A. Gallego

**Affiliations:** Department of Bioengineering, Imperial College London, London, UK; Département de neurosciences, Université de Montréal, Montréal, Canada; Department of Neuroscience, Northwestern University, USA; Department of Biomedical Engineering, Northwestern University, Evanston, IL, USA; Department of Physical Medicine and Rehabilitation, Northwestern University, and Shirley Ryan Ability Lab, Chicago, IL, USA

## Abstract

Animals use afferent feedback to rapidly correct ongoing movements in the presence of a perturbation. Repeated exposure to a predictable perturbation leads to behavioural adaptation that counteracts its effects. Primary motor cortex (M1) is intimately involved in both processes, integrating inputs from various sensorimotor brain regions to update the motor output. Here, we investigate whether feedback-based motor control and motor adaptation may share a common implementation in M1 circuits. We trained a recurrent neural network to control its own output through an error feedback signal, which allowed it to recover rapidly from external perturbations. Implementing a biologically plausible plasticity rule based on this same feedback signal also enabled the network to learn to counteract persistent perturbations through a trial-by-trial process, in a manner that reproduced several key aspects of human adaptation. Moreover, the resultant network activity changes were also present in neural population recordings from monkey M1. Online movement correction and longer-term motor adaptation may thus share a common implementation in neural circuits.

## Introduction

Animals, including humans, have a remarkable ability to rapidly correct their ongoing movements based on perceived errors, even when feedback may be distorted, such as when reaching into a pond to recover an object one has dropped. In the laboratory, these movement corrections and subsequent adaptation can be evoked and studied systematically using the classic visuomotor rotation (VR) perturbation paradigm^1, 2^. In this paradigm, the subject receives distorted visual feedback (typically a rotation about the centre of the workspace) of a reaching movement, thereby creating a perceived error due to the mismatch of expected and observed hand trajectory. Humans can correct their ongoing movements even during the very first trial after perturbation onset^3^, a process that is mediated by primary motor cortex (M1) integrating multiple feedback signals arriving from various sensory and motor brain regions^4–17^.

When repeatedly exposed to a predictable perturbation, animals progressively learn to use their perceived errors to anticipate its effect. For the case of the VR paradigm described above, this leads to a gradual reaiming of the reach, until it starts out in the correct direction^2^, thereby eliminating the need for further online corrections. This requires some form of rapid learning along the sensorimotor pathways, likely guided by trial-by-trial error information^3, 18, 19^. How and where in the brain this “motor adaptation” happens remains inconclusive^1, 20–45^, and may depend on the characteristics of the perturbation, such as whether it is a rotation of the visual feedback or a force acting on the limb^28, 46–51^.

Feedback-based movement correction and motor adaptation have been mostly studied in isolation and are often assumed to involve different neural substrates^2, 11^. What if that were not the case, but instead, they shared a common implementation in neural circuitry? A recent behavioural study proposed that movement correction and adaptation may indeed be tightly linked: fast feedback responses could act as teacher signals that drive trial-by-trial adaptation^21, 52, 53^. Despite the conceptual beauty of this idea, its feasibility and implementation details remain unexplored.

Here, we hypothesised that the neural circuitry for feedback control may be exploited to drive the “plasticity” that enables successful motor adaptation. To address this hypothesis, we use a recurrent neural network (RNN) model^54–62^ trained not only to produce a certain output, but to *control* it. The key difference here is that this model should be able to flexibly correct its output in the case of unexpected external perturbations. Having such a model allowed us to test whether feedback signals used for motor control could guide plastic changes within the network that lead to successful trial-by-trial learning.

We first show that an RNN can be trained to perform feedback-based motor control, even in the presence of a relatively long, biologically plausible feedback delay. We then demonstrate how feedback signals, modelled as inputs to the RNN, can guide synaptic plasticity within the network that drives successful trial-by-trial adaptation. Intriguingly, this form of learning through feedback-driven plasticity led to behavioural adaptation that was similar to that of a human in terms of time course^63, 64^, generalisation^2^, and sensitivity to perturbation variability^65, 66^. Moreover, both effective control and learning could be achieved with sparse feedback signals. Finally, comparison with neural population recordings from monkey M1 (data from^51^) supports the plausibility of the proposed plasticity rule: the temporally dissociable activity changes that followed adaptation in our model could also be found in the actual neural activity. This work not only introduces the potential of a combined implementation of motor control and learning in recurrent circuits, but it also relates several aspects of human adaptation behaviour to a single unifying neural process.

## Results

### A recurrent neural network that performs feedback-based motor control

We used an RNN model to investigate whether the same feedback signals used to control ongoing behaviour could also mediate motor adaptation. This work was divided into two phases: 1) training the RNN to perform feedback-based motor control (using a gradient-based algorithm), and 2) using this trained RNN to implement trial-by-trial motor adaptation via a local, biologically plausible learning rule acting on the recurrent weights of the network. We validated our model by comparing it to several behavioural and electrophysiological studies.

First, we trained an RNN to control its own output to produce the desired movement (Figure 1A,B). Our goal was to have a model that, after training, could dynamically adjust the ongoing movement according to incoming sensory feedback. Feedback control was based on an instantaneous position error signal, *ϵ*_*t*_, defined as the difference between the produced position, *p*_*t*_, and the target position, 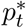, which we computed assuming a straight line between the start and end points (Figure 1C). This error signal was fed back as an input to the RNN with a biologically realistic delay of 120 ms^12^ (Figure 1B).

**Figure 1:**
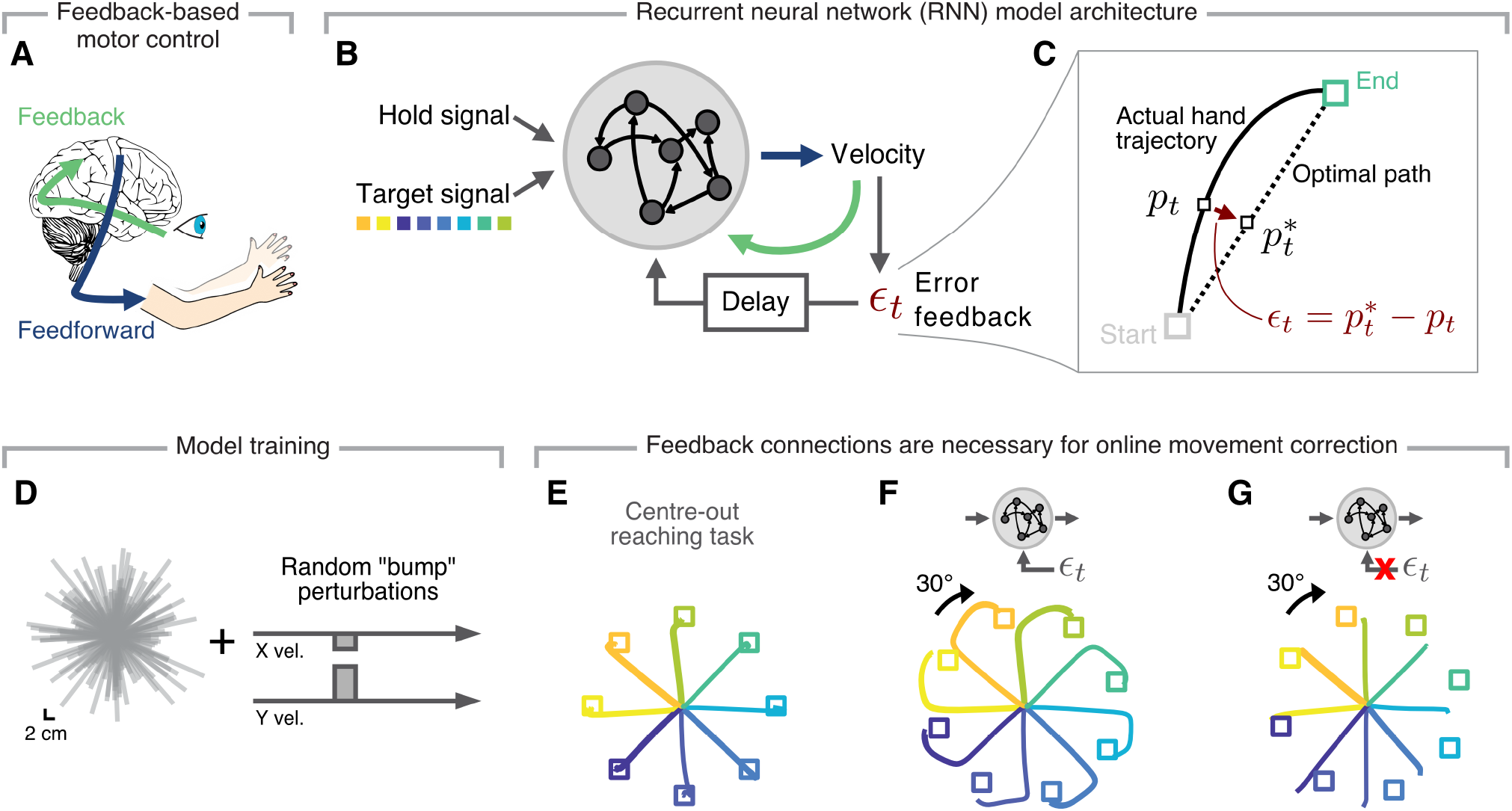
Proposed recurrent neural network that controls its output based on feedback. **A**. Afferent and efferent pathways act together to produce precise movements, and to flexibly correct them in the presence of perturbations. **B**. A recurrent neural network (RNN) model to explore a shared implementation of motor control and adaptation based on a common error feedback loop. **C**. We defined the ongoing error during movement as the difference between the observed and the optimal hand position. **D**. Initial RNN training included reaches of varying lengths and to different locations (grey lines) with occasional random velocity “bump” perturbations (cf. Figure S1). **E**. Hand trajectories produced by a trained RNN required to perform a standard eight-target centre-out reaching task. **F**. Hand trajectories after introducing a 30° rotation of the RNN’s output, to mimic a visuomotor rotation perturbation; note that feedback allowed the network to correct its output online and reach to the target. **G**. Hand trajectories for a model without a feedback loop, which could not counteract the same perturbation.

After the initial training phase (Figure 1D, examples in Figure S1; Methods), we tested the RNN on a standard centre-out reaching task with eight equally distributed targets. As expected due to our training procedure, the model was readily able to produce the required straight movement trajectories even without explicit training on this task (Figure 1E).

To test the network’s ability flexibly correct its output, we replicated the classic VR paradigm^2^. If the RNN could indeed *control* its output, it should still be able to reach to the desired target by correcting the produced movement to counteract the 30° rotations online. Inspecting the movement trajectories after VR onset confirmed that the model could use the error signal to correct its ongoing output (Figure 1F; the curved trajectories indicate ongoing correction). Importantly, successful correction relied on the error signal being fed back into the model: trajectory correction only started after the delayed feedback had had enough time to propagate to the model (Figure 1F), and an RNN trained without feedback connections (Methods) could not reach to the targets (Figure 1G).

### An error feedback signal used for feedback-driven motor control can drive trial-by-trial adaptation

We have shown that an RNN that has learnt to use feedback signals to control its output can readily counteract an external perturbation, exhibiting a behaviour upon VR onset that is very similar to that of humans^2^ and monkeys (compare the monkey data from Ref. 51 in Figure 2A to the model data in Figure 2D). However, when repeatedly exposed to a VR, both humans and monkeys learn to adjust their initial “motor plan,” which results in their reach take-off angle pointing in the correct direction after dozens of trials^2, 51^ (Figure 2B,C). Can our relatively simple error feedback signal enable persistent learning across trials?

**Figure 2:**
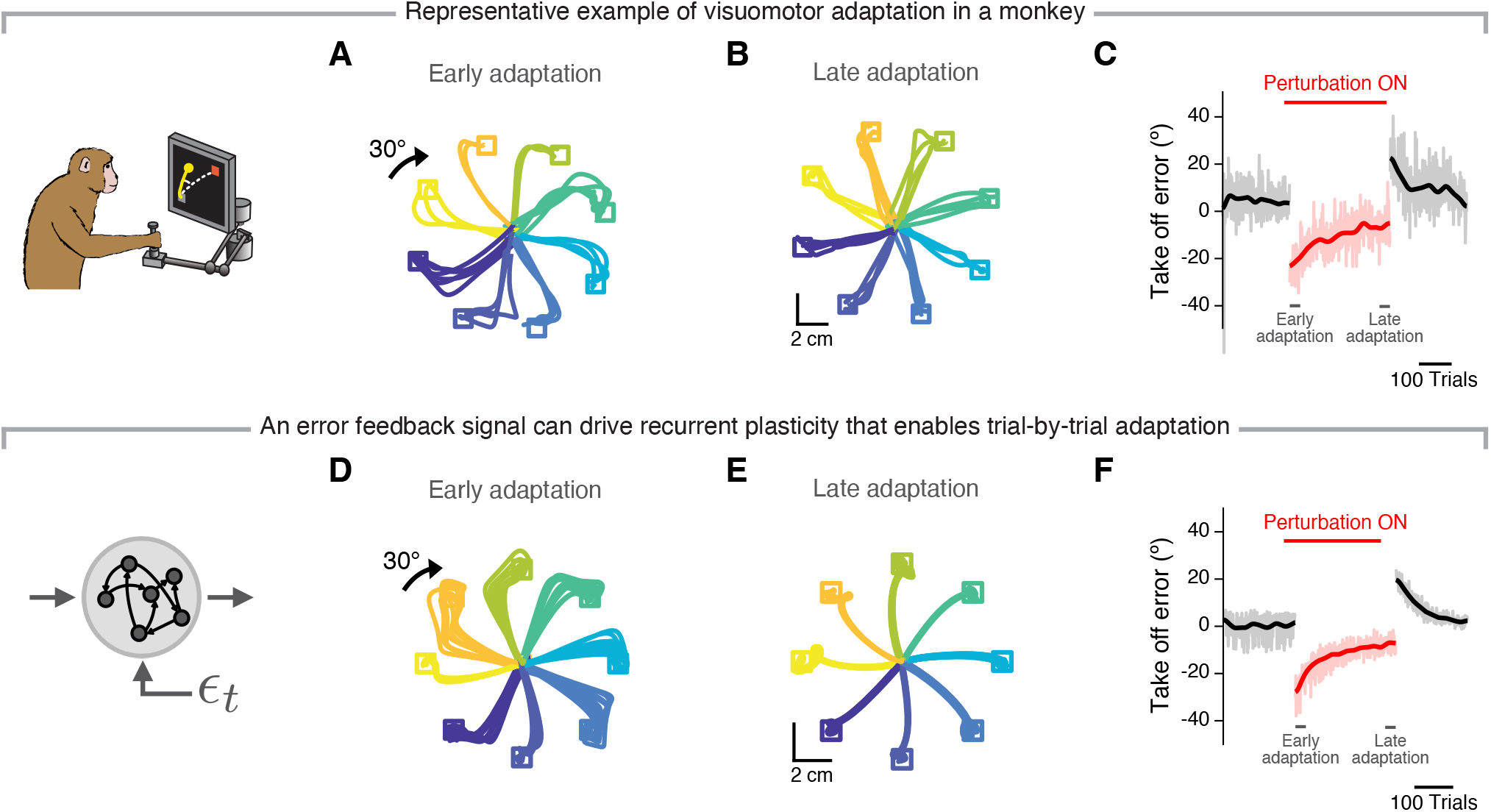
Feedback signals can guide local synaptic plasticity that enables successful motor adaptation. **A**. Example hand trajectories as a monkey reached to each of eight targets after a 30° rotation of the visual feedback was introduced (first 30 trials after perturbation onset; data from Perich et al.^51^). **B**. Example hand trajectories after the monkey had adapted to the perturbation by reaiming the reach direction (last 30 trials of the perturbation phase for the same session as in A). **C**. Angular reaching take-off error for the baseline (left, black), perturbation (middle, red), and wash-out phases (right, black). Transparent lines, single trial errors; solid lines, smoothed mean error (Gaussian filter, s.d., 10 trials). **D-F**. Simulation results for an RNN implementing the proposed feedback-driven plasticity rule, by which recurrent weights are modified according to the error signal received by the postsynaptic neuron. The first and last 80 trials of adaptation are shown; otherwise data is presented as in A-C. Note the strong similarities between the behaviour of the network and that of the monkey.

Since the feedback inputs acting on the network correctly modulate each neuron’s activity to minimize the ongoing motor error (Figure 1F), we hypothesised that they could also act as a “teacher signal” for local, recurrent synaptic plasticity. To test this, we devised a local biologically-plausible synaptic plasticity rule causing the connection weight from neuron *i* to neuron *j* to change in proportion to the feedback signal received by neuron *j* (Methods). Implementing this plasticity rule led to behaviour that was similar to that of monkeys’: the initially large errors in take-off angle became progressively smaller over time, until they reached a plateau close to zero error (note the similarities between Figure 2D-F and Figure 2A-C). Moreover, when the perturbation was turned off, the model underwent a de-adaptation phase similar to the “wash-out” effect exhibited by monkeys (compare Figure 2C and Figure 2F) and humans^67^. These results confirm our main hypothesis: an error feedback signal used for online motor control can be leveraged to guide recurrent synaptic plasticity that drives successful trial-by-trial motor adaptation.

### Feedback-based learning recapitulates key features of human motor adaptation

The previous simulation results suggest that a relatively simple plasticity rule based on an error feedback signal may mediate motor adaptation. Could this type of learning be implemented in actual brains? To investigate the biological plausibility of our feedback-based plasticity rule, we tested whether our model replicated various key observations from human and monkey motor adaptation studies.

We first addressed the finding that humans learn more from a given trial if they experience a larger error^52^. Indeed, our model reproduced this trend; the measured correlations between movement error and amount of learning in the next trial were comparable in magnitude and sign to those of monkeys performing the same VR task (Figure 3A; data from Ref. 51). Moreover, as in human adaptation studies^65, 66^, our model’s ability to learn was also hindered when the perturbation was inconsistent across trials, with greater perturbation variance leading to progressively less learning (Figure 3B). Thus, the amount of trial-by-trial adaptation matched experimental observations well.

**Figure 3:**
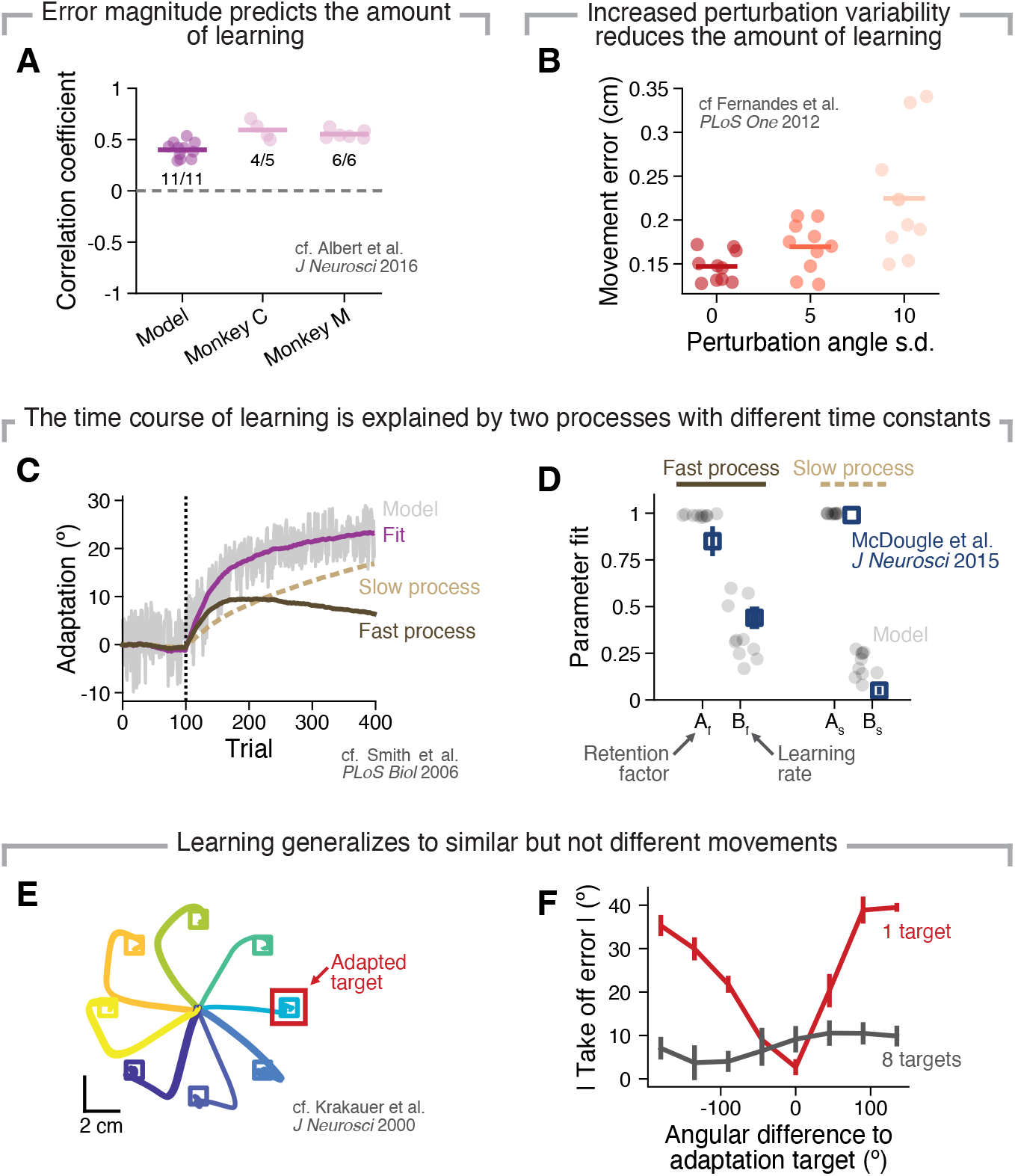
Motor adaptation based on feedback-driven plasticity recapitulates key aspects of human and monkey behaviour. **A**. Correlation between the take-off error in the current trial and the amount of learning from this trial to the next for both simulation (Model) and behavioural data (Monkey C and Monkey M; data from Perich et al.^51^). Individual circles, a different network (Model) or experimental session (Monkey); numbers, proportion of networks or experimental sessions exhibiting a significant correlation (*P<*0.05). **B**. Movement error after adaptation to VR perturbations with different perturbation angle variability (cf. Ref. 65, 66 for human behaviour). **C**. As in human experiments, the time course of adaptation in the model is well fitted by a dual-rate model^63^. Grey line, single trial errors of simulated adaptation behaviour in an example simulation. Purple line, model fit; dark brown line, fast process; light brown line, slow process. **D**. Fitted parameters of the dual rate model (black) match those from a visuomotor adaptation study in humans^64^ (blue). Individual circles, ten different networks; square and error bars, mean and 95 % confidence interval. **E**. Hand trajectories produced by the model after visuomotor adaptation to a single target (“Adaptation target”, in red). **F**. Take-off error after adaptation to visuomotor perturbations applied to a single target, as shown in E (red), or to eight targets (dark grey) (cf. Ref. 2 for human behaviour). Lines and error bars, mean and s.d. across ten networks.

When examining the timescale of learning during an experimental session, human motor adaptation seems to be mediated by two simultaneous learning processes: one fast, and another slow^63, 64^. Using the same analysis as in Ref. 63, we found that the adaptation time course of our model is also best described by a combination of two learning processes with different time constants (Figure 3C). Even the parameters describing these processes were comparable to values reported in a human VR adaptation study (Figure 3D; data from Ref. 64), indicating that, as in actual experiments, our feedback-driven plasticity rule is dominated by fast learning early in adaptation and slow learning later on.

Finally, animals, including humans, have a remarkable ability to generalise what they have learned to novel situations, yet, the amount of generalisation seems to depend on the similarity between the current and the past situation. During a VR experiment, participants who have adapted to a perturbation applied on a single reach direction generalize when reaching to neighbouring targets, to an extent that decreases as the angle between the new and adapted direction increases^2^. Repeating this single target adaptation experiment in our model revealed the same kind of generalization: the model readily anticipated perturbations applied on adjacent targets and adaptation decreased as the angle between the probed target and the adapted target increased (Figure 3E-F). In summary, our model reproduced key features of primate motor adaptation, supporting our unified view of how rapid motor learning could be implemented in the brain by leveraging signals mediating online feedback correction.

### Sparse feedback is sufficient for both motor control and adaptation

The fact that our model recapitulates many aspects of human and monkey adaptation behaviour supports the hypothesis that a similar feedback-based learning process may be implemented in actual neural circuits. Yet, in the model we have examined so far, every unit received a feedback signal, whereas only a relatively large subset of primary motor cortical neurons seem to be modulated by sensory feedback (73% of neurons in M1 respond to either visual or proprioceptive feedback^16^). Thus, to further probe the plausibility of our model, we explored how our results were influenced by feedback connection density.

We quantified model performance in terms of both control accuracy and degree of learning for feedback projection densities ranging from 1% to 100%. Critically, the model could control its output effectively with density as low as 25%; even the extremely sparse 1% projection density decreased endpoint error considerably compared to a model without feedback (compare the solid and dashed lines in Figure 4A). Likewise, the model did learn substantially for all feedback projection densities (compare the solid and empty markers in Figure 4B), and, although the amount of learning decreased almost linearly as feedback projections became sparser, as few as 1% of the neurons receiving feedback sufficed to drive meaningful learning that reduced take-off errors by 30%. Notably, these results were robust to changes in RNN connectivity, as control accuracy and degree of learning were similar for recurrent connection probabilities of 50% and 80% (Figure S2).

**Figure 4:**
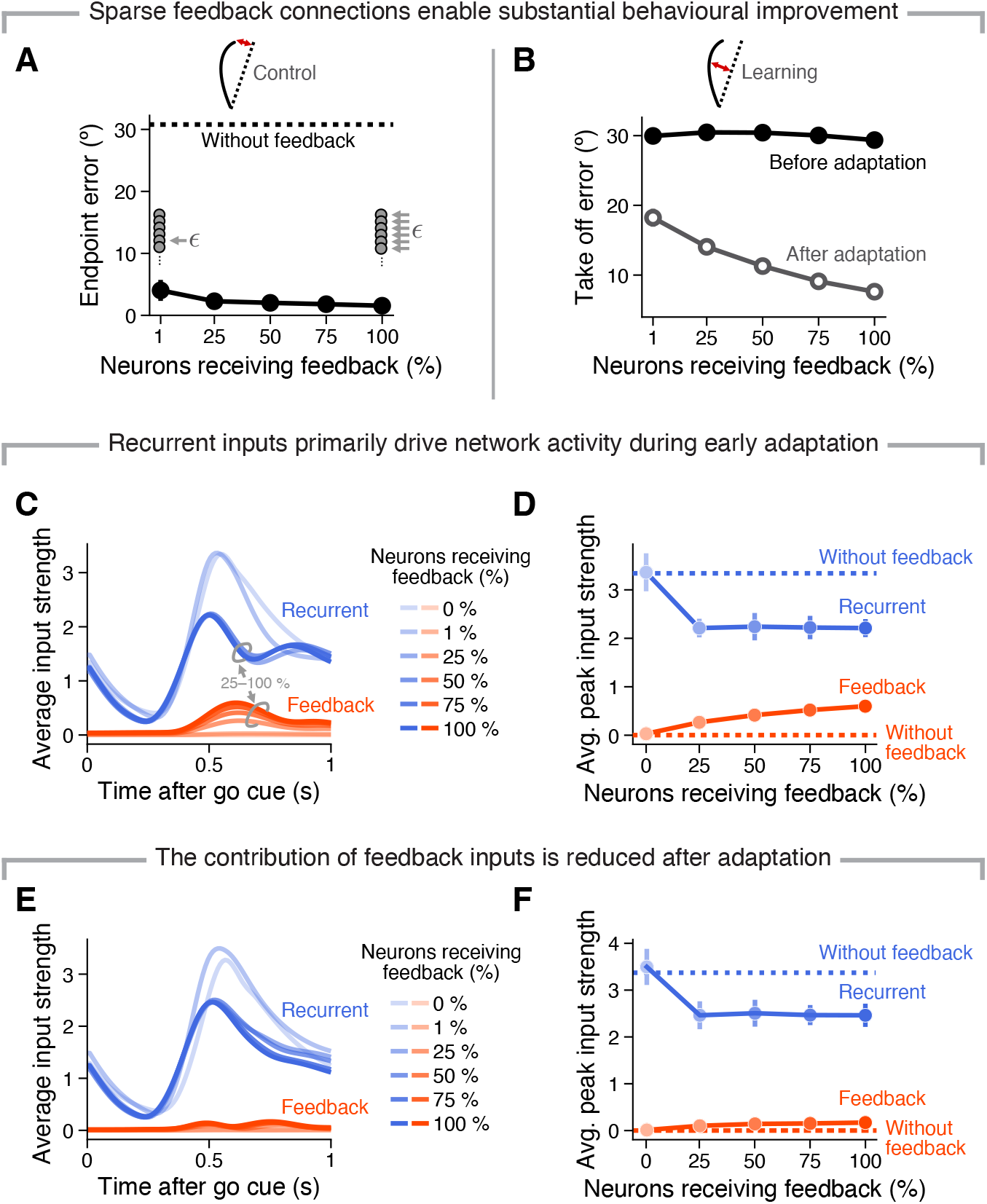
Both feedback-based motor control and adaptation can be achieved with sparse feedback signals which are small compared to recurrent signals. **A**. Angular error between produced and target position at the end of the reach immediately after onset of visuomotor rotation for networks with different percentages of neurons receiving afferent feedback (black markers), including no feedback (dashed line). Lines and error bars, mean and s.d. across ten networks. **B**. Take-off error at visuomotor rotation onset (solid circles), and after adaptation (empty circles) for networks with different percentages of neurons receiving afferent feedback. Lines and error bars, mean and s.d. across ten networks. **C**. Average recurrent (blue) and feedback (red) inputs to an RNN neuron before adaptation. Average input strength is defined as the mean across incoming signals and neurons. Legend, percentage of neurons receiving feedback. **D**. Average magnitude of the peak input strength of the recurrent (blue) and feedback (red) inputs before adaptation. Same colour scheme as in C. Lines and error bars, mean and s.d. across ten networks. **E**. Average recurrent (blue) and feedback (red) inputs to an RNN neuron after adaptation. Data presented as in C. **F**. Average magnitude of the peak input strength of the recurrent (blue) and feedback (red) inputs after adaptation. Data presented as in D.

Finally, to understand how feedback inputs enabled successful behaviour, we compared their contribution to that of the recurrent inputs, both early in the perturbation block (Figure 4C,D), where feedback is crucial to correct the output, and after successful adaptation (Figure 4E,F), where we expected feedback to only play a minor role. Even at perturbation onset, the network activity was mostly driven by recurrent inputs, with the feedback making up only a minor part of the overall input (compare the blue and red traces in Figure 4C,D, respectively); this was the case for all feedback densities. Thus, feedback signals did not directly drive accurate movements; instead their contribution was amplified through recurrent dynamics —note the different temporal patterns of recurrent input between before and after adaptation (Figure 4C and Figure 4E, respectively). This likely explains why movement accuracy remained stable across a broad range of feedback projection densities (Figure 4A). As expected, after successful adaptation, feedback inputs were much smaller than recurrent inputs, regardless of the feedback projection density (Figure 4E,F). Combined, these results suggest that actual brains may perform effective motor control and adaptation even with a relatively small number of feedback connections, and feedback signals may make relatively small contribution to the overall neural activity.

### Two temporally dissociable adaptation-related activity changes in the model can be uncovered in monkey primary motor cortex

Our previous simulations suggest that sparse feedback projections may suffice to enable effective motor control and adaptation (Figure 4A,B), and that feedback inputs account for only a minor portion of a unit’s overall input (Figure 4C,D). Combined, these observations indicate that identifying a “signature” of feedback-driven plasticity in neural recordings or even in the RNN activity may be difficult. Our approach to tackle this was to focus on the within-trial timing: we devised an analysis to isolate activity changes related to the two temporally distinct processes that we expected to occur during feedback-driven adaptation: 1) an early feedforward change, reflecting updated pre-planned movement intent, and 2) a change in feedback signals later in the trial, as the error decreased over trial-by-trial learning.

To measure feedback related changes (Figure 5A; green) we focused on the activity changes from the baseline epoch (box A in Figure 5A) to right after perturbation onset (box B in Figure 5A), by simply taking the difference between the single neuron activities in those two epochs. Similarly, the learning related changes (Figure 5A; blue) represent the activity changes between the baseline epoch and the late adaptation epoch (box C in Figure 5A), where we expect the feedback component to have reached baseline levels again as the model no longer needs to correct its movement online. The adaptation related change (Figure 5A; dark grey) is defined as the difference between early and late adaptation epochs, respectively.

**Figure 5:**
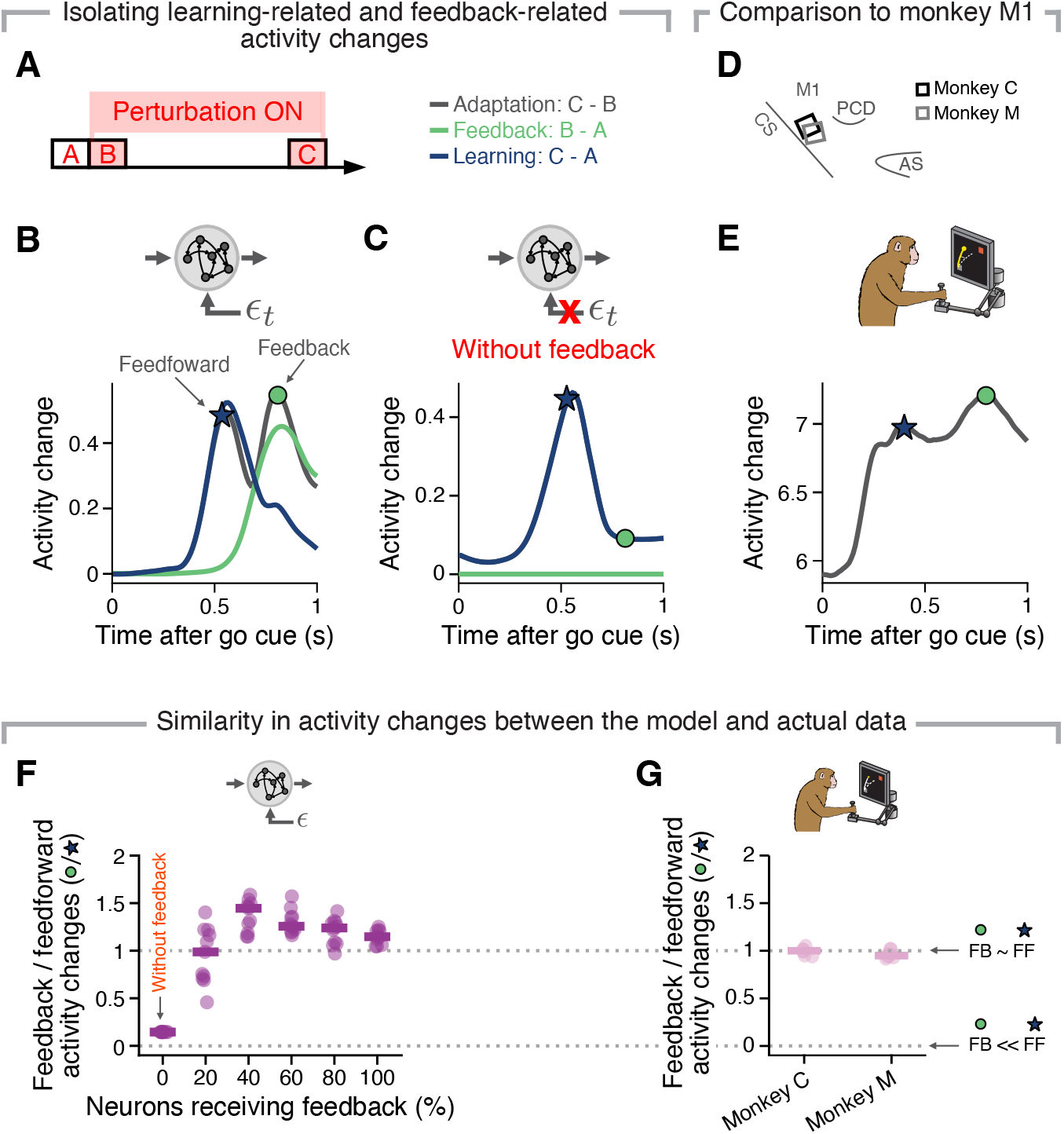
Two temporally dissociated activity changes may indicate feedback-driven plasticity in monkey M1 during visuomotor adaptation. **A**. Epochs used to identify feedback-related (green) and learning-related (blue) activity changes. **B**. The average RNN activity change across pairs of behavioural epochs (as shown in A) reveals putative feedforward-related (blue star) and feedback-related (green circle) activity changes for an example network. Black trace, activity change between perturbation onset and successful adaptation; green, activity change between baseline and perturbation onset; blue, activity change between baseline and successful adaptation. Note that the peak timings are chosen based on ten different network simulations. **C**. An RNN without a feedback loop has only feedforward-related activity changes. Data presented as in A. **D**. Approximate location of the recording array for each of monkey (legend). **E**. The average activity change of monkey M1 neurons between perturbation onset and successful adaptation resembles that of the network (compare with A). Data from one representative session from Monkey C, presented as in B. **F**. Comparison between the ratio of the feedback-related (green circle in B,C,E) to feedforward-related (blue star in B,C,E) activity changes from perturbation onset until successful adaptation for networks with different feedback densities. Individual markers, individual networks. Horizontal line, mean. **G**. Same as F, but for monkey M1 (data shown for each monkey separately). Individual markers, individual sessions. Horizontal line, mean. FB, feedback; FF, feedforward.

Inspecting the RNN activity changes during adaptation indeed revealed two distinct peaks in the average activity change during adaptation (Figure 5B), assigned to learning (blue) and feedback (green), respectively. To confirm that these early and late “activity change peaks” did reflect feedforward and feedback processes, respectively, we trained a different set of models that used simple gradient descent instead of feedback-driven plasticity to achieve motor adaptation (Methods). As expected, these models also exhibited a change in activity early during the movement (compare the blue traces in Figure 5B and Figure 5C), confirming that it does represent learning. In contrast, the second peak (green trace in Figure 5B) was absent from these models without feedback-driven plasticity, confirming its source.

Having uncovered a signature of feedback-driven adaptation in our model, we sought to identify a similar change in neural population recordings from monkey M1 (Figure 5D; data from Ref. 51; Methods). Figure 5E shows the average change in M1 activity during a representative VR adaptation session, qualitatively confirming our prediction of a feedback signal during early adaptation trials (circle in Figure 5E). Reassuringly, this feedback signal occurred 300 ms after peak speed (which happened 500 ms after the go cue), as it did in other studies exploring feedback-based movement corrections^12, 16^. To further compare the monkey data to our model, we calculated the ratio of the adaptation-related activity change at the “feedback peak” (green circle in Figure 5B,C,E) and the learning-related “feedforward peak” (blue star in Figure 5B,C,E) (Methods). For both monkeys, there was a substantial change in neural activity at the feedback peak during adaptation (Figure 5G), as predicted by our model (Figure 5F). Moreover, the value of this ratio was similar across both monkeys and sessions, and best matched, in terms of mean and variability, that of models with high feedback projection densities (Figure 5F,G). The fact that models with dense feedback projections best replicated the experimental data is in good agreement with observations suggesting that as many as 73% of M1 neurons are driven by feedback^16^. The similarities in the activity changes during adaptation between our model and monkey M1 suggest that error feedback signals in M1 may indeed guide local plasticity relevant for rapid motor learning.

## Discussion

Both the rapid correction of ongoing movements and progressive adaptation to changing conditions are key landmarks of behaviour that are often studied separately. Here, we have shown that an RNN that can dynamically control its output based on feedback inputs can use those same inputs to achieve motor adaptation through recurrent connectivity changes. Interestingly, this form of feedback-driven plasticity recapitulated key aspects of primate adaptation behaviour^2, 63–66^, and led to identifiable activity changes in the model that we subsequently discovered in neural recordings made in the primary motor cortex of two monkeys. These results, which were robust across a broad change of model parameters (Figure 4, Figure S2), support the hypothesis that online movement correction and motor adaptation may share a common implementation in neural circuits.

### Implementation of feedback-based motor control in neural circuitry

Most modelling studies on motor control have focused on understanding it at an abstract computational level, mostly ignoring neurons and connections between them^52, 63, 68^. One potential reason is the challenge of mapping the abstract concepts of optimal feedback control theory into brain regions^5, 17^. Our work differs from those previous attempts in that it approaches the problem from a bottom-up, not a top-down perspective: instead of explicitly training neural networks to match the behavioural components predicted by optimal feedback control theory, such as a forward model or a controller^69^, we let error minimization guide the emergence of an efficient control strategy, potentially mimicking how brain connectivity developed over evolutionary timescales^70–72^. Given that this bottom-up approach led to a feedback-based learning process that recapitulated key aspects of human behavioural adaptation (Figure 2,Figure 3), we believe that our model could help map specific functions that have been formalized in optimal feedback control onto neural circuits, adding to existing approaches^15, 17, 73–76^.

While in this paper we have compared the network activity to recordings from monkey M1, our model (Figure 5) does not necessarily encompass only M1 function. On the contrary, it is likely that it captures functions mediated in part through multi-region interactions with a variety of cortical and subcortical regions^77, 78^, such as feedback processing, or sensory gating. Extending our work to modular, multi-region RNNs whose activity is compared to neural population recordings across the sensorimotor network (such as in Ref. 59, 61, 62) may shed light into the distributed implementation across brain circuits of the different computations underlying feedback control.

Early studies proposed the cerebellum as the key structure for motor adaptation^5, 22, 25, 31, 41, 79–84^. Their central premise was that the cerebellum stores both “forward” and “inverse” internal models of the sensorimotor system which are used to generate and update predicted outcomes of motor commands, respectively^50, 85–87^. The importance of internal models in the cerebellum received additional support from evidence of adaptation deficits in cerebellar patients^88^, although this view has recently been challenged by a proposal that adaptation is better described by a direct update of a control policy^53^. Our study does not contradict this work; it illustrates an alternative, perhaps parallel learning process relevant for motor adaptation. Moreover, the cerebellum could readily fit into our model as the brain region responsible for calculating error estimates based on the sensory feedback, which we simply assumed to exist. Interestingly, the fact that cerebellar patients have impairments in motor control as well as learning^25, 89^ also fits with our model, since both processes are critically dependent on the availability of an accurate error signal. Combining our bottom-up model of feedback-based learning in M1 with previous models of the cerebellum^22, 76^ might provide a fruitful route to uncover how each of these two brain regions contributes to motor adaptation.

### Biologically plausible learning rules

Our proposed feedback-driven learning rule adds to recent efforts to implement biologically plausible learning in RNNs^90–96^ —a challenging temporal credit assignment problem. Most of these learning rules share the same basic idea: the weight change is proportional to the error signal arriving at the postsynaptic neuron times the activity history of the presynaptic neuron. What makes those learning rules biologically plausible is that both pieces of information could in principle be locally available at a synapse.

The crucial difference between our model and previous studies is that the error signal is a direct input to the neuron, which allows it to simultaneously guide weight update and affect the ongoing network dynamics. This feature is desirable because it avoids the need to have two distinct pathways for error signals and ongoing network dynamics, respectively^97–99^. Moreover, our approach makes weight update dependent on the error signal in two ways: directly, through the error term in the plasticity rule, and indirectly, through the change in ongoing dynamics, which influences the activity of the presynaptic neuron. This could be beneficial for learning and should be compared to standard learning algorithms like gradient descent.

Finally, our feedback-driven plasticity rule led to an adaptation time course that replicated the simultaneous fast and slow processes found in human behavioural studies (Figure 3C;^63, 64^). This seems puzzling given that the fast process is often assumed to be explicit^64, 100^, a component not possible in our RNN implementation. Therefore, the observation of a fast component despite adaptation being mediated purely by an error feedback signal suggests that, in principle, error minimisation alone can give rise to learning at multiple timescales.

## Conclusions

We have shown how the same error feedback signal that mediates ongoing motor corrections can drive trial-by-trial motor adaptation by guiding synaptic plasticity using a relatively simple, biologically plausible plasticity rule. Crucially, this bottom-up model of a shared implementation of feedback-based motor control and rapid learning recapitulated key observations from behavioural and neurophysiological adaptation studies in humans and monkeys. Thus, these two landmarks of animal behaviour may be unified in their implementation by the same neural circuitry.

## Methods

### Recurrent Neural Network model

Neural activity *x* was simulated using the following dynamical equations,

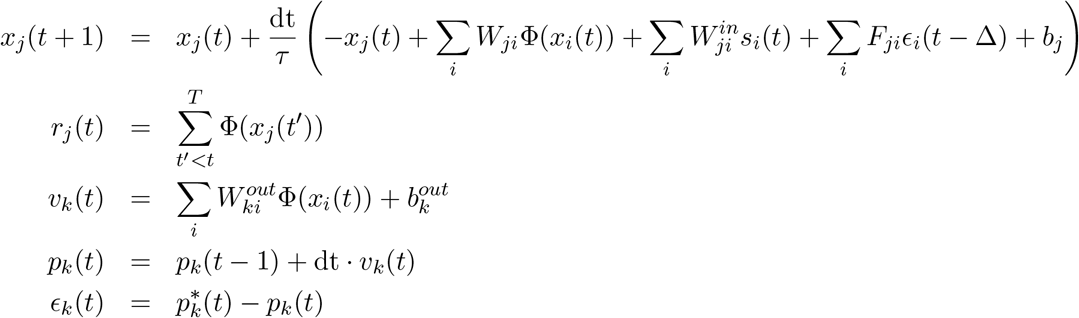

where the network output *v* represents velocity, and *p* position, of the simulated planar hand movement (cf. definitions in Table 1). The instantaneous error signal *ϵ* is given by the difference between the target *p*^*^ and the produced position *p*, and is fed back to the network with a time delay Δ. How we constructed the network input *s* and the target position *p*^*^ are described in the “Reaching datasets for model training and testing” section below. Each trial was initialized by setting all *x*_*j*_ to random numbers uniformly distributed between -0.2 and 0.2. All simulations were performed on RNN models consisting of 400 neurons, which were connected all-to-all. Varying the recurrent connection probability did not change the results (Figure S2).

**Table 1:** Simulation parameters.

| Parameter | Definition | Value |
| --- | --- | --- |
| $\text{dt}$ | Time step | 10ms |
| $\tau$ | Time constant | 50ms |
| $\Delta$ | Feedback delay | 120ms |
| $N$ | Number of neurons | 400 |
| $\Phi$ | Nonlinearity | ReLU |
| $\Phi(x)$ | Neural activity | - |
| $s$ | Stimulus | - |
| $\epsilon$ | Position error | - |
| $r$ | Eligibility trace | - |
| $W$ | Recurrent weight matrix | - |
| $b$ | Recurrent offset | - |
| $W^{in}$ | Input weight matrix | - |
| $W^{out}$ | Output weight matrix | - |
| $b^{out}$ | Output offset | - |
| $F$ | Feedback weight matrix | - |
| $v$ | Velocity (2D) | - |
| $p$ | Position (2D) | - |
| $\alpha$ | Gradient Descent: Learning rate | 0.001 |
| $B$ | Gradient Descent: batch size | 20 |
| $\beta$ | Gradient Descent: weight regularization | 0.001 |
| $\gamma$ | Gradient Descent: activity regularization | 0.002 |
| $\eta$ | Feedback-driven plasticity: learning rate | 0.000001 |
| $T$ | Reach trajectories: number of time steps in a trial | 300 |

### Model training procedure

The first step was to train the RNN to control its own output, that is, to minimize the position error, *ϵ*. This training was performed using standard gradient descent to find the right set of parameters. The initial training procedure was implemented in Pytorch^101^ using the Adam optimizer with learning rate *α* = 0.001 (*β*_1_ = 0.9, *β*_2_ = 0.999)^102^. The weights (*W, W*^*in*^, *W*^*out*^, *F*) and biases (*b, b*^*out*^) were initialized by drawing random, uniformly distributed numbers between 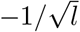 and 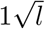 where *l* is either the number of neurons in the network (for *W, W*^*in*^, *F, b*) or the dimensionality of the output (for *W*^*out*^, *b*^*out*^). The gradient norm was clipped at 0.2 prior to the optimization step. The loss function used for this initial training phase was defined as

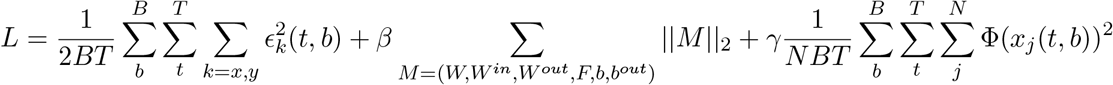

where *B* is the batch size, *T* the number of time steps, *N* the number of neurons, *β* the regularization parameter for the weights and bias terms and *γ* the regularization parameter for the activity in the network (cf. definitions in Table 1). The network was trained for 1100 epochs, divided into three blocks of different lengths (100, 500, 500). For the first 100 epochs, the feedback weights *F* were kept fixed while the remaining parameters where allowed to change. This ensured that the model learnt to self-generate the appropriate network dynamics to produce a variety of reaching trajectories. In the next 500 epochs, the feedback connections were also allowed to change. In the last 500 epochs, we introduced perturbations on the produced output (see “Reaching datasets for model training and testing”), with all parameters plastic, to make the model learn to use the feedback inputs to compensate for ongoing errors.

### A feedback-driven plasticity rule to drive trial-by-trial learning

Having set up the model to control its own output, we next examined how feedback inputs *ϵ* could guide learning, implemented through synaptic plasticity within the recurrent weights *W* of the network. To this end, we devised the following *feedback-driven plasticity rule*:

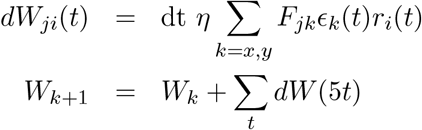

The weight update *dW* was calculated online and summed up taking into account every fifth time step until the end of a trial *k*. After each trial, we applied this accumulated weight change and updated the recurrent weights *W* accordingly.

### Reaching datasets for model training and testing

The network model was trained to produce a broad set of synthetic planar reaching trajectories following an instructed delay phase. The *x* and *y* positions of the starting (*p*^*start*^) and ending points (*p*^*end*^) of those trajectories were randomly drawn from a uniform distribution ranging from -6 cm to 6 cm. To simulate natural reaching behaviour, we interpolated between these points using a sigmoid function

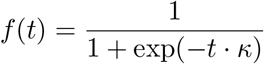

where *κ*=10/s. The manually constructed reach trajectories were thus given by

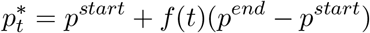

which resulted in bell-shaped velocity profiles.

Each trial lasted 3 s and included an instructed delay period, randomly drawn from 0 s to 1.5 s. The network received an input signal consisting of a two-dimensional target signal and a one-dimensional timing signal. The target signal was defined as (*p*^*end*^ − *p*^*start*^). It was delivered to the network 0.2 s after trial onset, and was fixed until the end of the trial. The timing signal was given in form of a constant, and was switched to zero at the time corresponding to the “go” signal, which varied between 0.2 s and 1.7 s for the random reaching task used during training (that is, when the network generated reaches of random direction and lengths up to 8.5cm), and between 1.2 s and 1.7 s for the centre-out-reaching task.

As mentioned above, during the last phase of the initial training phase we included brief “bump” perturbations to the output of the network so it had to learn to use the feedback input to correct its output online. In 75 % of the trials, we added a pulse of 0.1 s duration and amplitude 10 cm/s on the velocity output of the model, either in *x* or *y* direction. This pulse occurred randomly between 0.2 s and 1.9 s after trial onset, to mimic perturbations at various movement periods.

After training, we tested the model on a centre-out-reaching task with eight targets equally distributed on a circle of 5 cm radius. In this task, the go signal occurred randomly between 1.2 s and 1.7 s, as before.

To probe online feedback correction and motor adaptation, we introduced a visuomotor rotation (VR) perturbation that rotated the output of the model by 30°, similar to previous visuomotor rotation experiments in humans^2^ and monkeys^51^.

### Neural recordings from behaving monkeys

We reanalysed previously published data from two macaque monkeys performing a visuomotor adaptation reaching task with a cursor controlled by movements of a manipulandum. In each session the monkeys performed 154–217 successful trials of an eight target centre-out reaching task. After this baseline period, a 30° rotation (clockwise or counter clockwise, depending on the session) of the cursor position feedback presented on a screen was introduced. Finally, after 219–316 successful adaptation trials, the perturbation was removed in order to study de-adaptation during this “washout” period.

We analysed the activity of populations of putative single neurons recorded using 96-channel microelectrode arrays chronically implanted in the arm area of primary motor cortex (details in Ref. 51). We quantified trial-by-trial learning by examining the monkey’s hand trajectories, which was tracked by recording the position of the handle of the manipulandum. All surgical and behavioural procedures were approved by the Institutional Animal Care and Use Committee at Northwestern University (Chicago, USA).

## Data analysis

### Movement error metrics to quantify learning

The take-off angle was defined as the initial reach direction, calculated between the go cue and peak velocity. When pooling the angular error across monkeys in Figure 2, we smoothed the mean across all sessions from both animals using a Gaussian filter with s.d. of ten trials.

When studying how error magnitude influences learning in the next trial (Figure 3A), we computed the Pearson’s correlation (*pearsonr* from `scipy.stats` package) between the absolute value of the angular error and the difference in angular error between the current trial and the next trial. To assess whether these correlations were significant, we compared them to a null distribution under the assumption of joint normality. The movement error in Figure 3B is defined as the averaged squared position error *ϵ*.

### Analysis of learning timescales

We investigated whether our model’s learning time course is composed of two processes with different timescales by implementing the analysis used in earlier studies Ref. 63, 64. We fitted a dual-rate state-space model to the angular error data, defined as below:

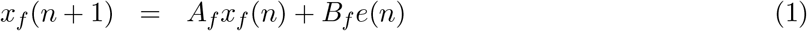

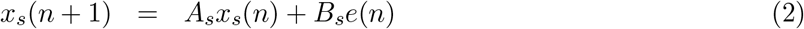

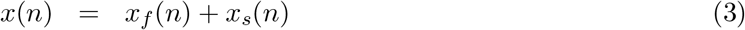

subject to the constraints *A*_*f*_ *< A*_*s*_, *B*_*f*_ *> B*_*s*_. *A*_*f*_ and *B*_*f*_ are the parameters describing the fast process, whereas *A*_*s*_ and *B*_*s*_ are the parameters describing the slow process. The *Adaptation* variable, *x* (cf. Figure 3C), is defined as the amount of change in the take-off reaching direction. The error *e* is given by the take-off angular error scaled to [-1,1]. The four parameters *A*_*f*_, *A*_*s*_, *B*_*f*_, *B*_*s*_ were obtained by fitting the model to the adaptation time course observed in the simulated data. For this, we used the *Sequential Least Squares Programming* method from the scipy optimization package.

### Analysis of temporally dissociable adaptation-related activity changes

We sought to identify a “neural signature” of adaptation-related activity changes in the network that could be seen in neural recordings from behaving monkeys. To this end, we probed three different “behavioural epochs” as follows (Figure 5A). For the model data, we simulated 200 baseline trials (epoch A in Figure 5A), 200 trials beginning immediately after perturbation onset (prior to any learning; epoch B), and 200 trials beginning 300 trials after the onset of learning (epoch C). For the monkey data, we considered the following: 100 baseline trials (epoch A), the first 100 trials after perturbation onset, during which monkeys were beginning to adapt (epoch B), and the last 100 perturbation trials, when monkeys had learned to counteract the perturbation (epoch C). Note that for the monkey data, the feedback epoch B was not as clearly defined as for the simulation data, since the monkeys had already started learning within epoch B.

The activity change in the simulation data was calculated by measuring, for each unit, the activity difference between all pairs of behavioural epochs (A, B, C in Figure 5A). For this, we simulated the same trials (using the same random seed) without perturbations (A), with perturbations (B), and with perturbations after the network had adapted (C). To identify the time point within a trial at which the largest activity change happened, we computed the absolute value of activity change, and averaged the respective differences across neurons and trials. This resulted in the time courses shown in Figure 5A,B. For the monkey data (Figure 5E), since we could not have the exact trials in different epochs, we calculated the difference between trials in different epochs in an all-to-all fashion, then averaged over those trial pairs.

After an initial analysis of the average activity change across all ten RNN models, we could define a “feedforward” time point (0.5 s after the go cue), in which the largest activity change between late adaptation (epoch C) and early adaptation (epoch B) happened, and a “feedback” time point (0.8 s after the go cue), in which the largest activity change between early adaptation (epoch B) and baseline (epoch A) happened. These values were very similar to those identified in the analysis of neural recordings from monkey M1: feedforward time point, 0.4 s after the go cue; feedback time point, 0.8 s after the go cue. For the pooled analysis presented in Figure 5F,G, we took the values of the activity change traces at those time points and calculated the ratio between the value at the feedforward time point and the value at the feedback time point.

## Data availability

The data that support the findings in this study are available from the corresponding authors upon reasonable request.

## Code availability

All code to reproduce the main simulation results will be made freely available upon publication on GitHub (https://github.com/babaf/feedback-driven-plasticity).

## Author contributions

B.F., C.C. and J.A.G. devised the project. M.G.P. and L.E.M. provided the monkey datasets. B.F. ran simulations, analysed data and generated figures. B.F., C.C. and J.A.G. interpreted the data. B.F., C.C. and J.A.G. wrote the manuscript. All authors discussed and edited the manuscript. C.C. and J.A.G. jointly supervised the work.

## Acknowledgements

L.E.M. received funding from the NIH National Institute of Neurological Disorders and Stroke (NS053603 and NS074044). C.C received funding from the BBSRC (BB/N013956/1 and BB/N019008/1), the EPSRC (EP/R035806/1), the Wellcome Trust (200790/Z/16/Z), and Simons Foundation (564408). J.A.G. received funding from the EPSRC (EP/T020970/1) and the European Research Council (ERC-2020-StG-949660). The funders had no role in study design, data collection and analysis, decision to publish, or preparation of the manuscript.

## Supplementary Figures

**Figure S1:**
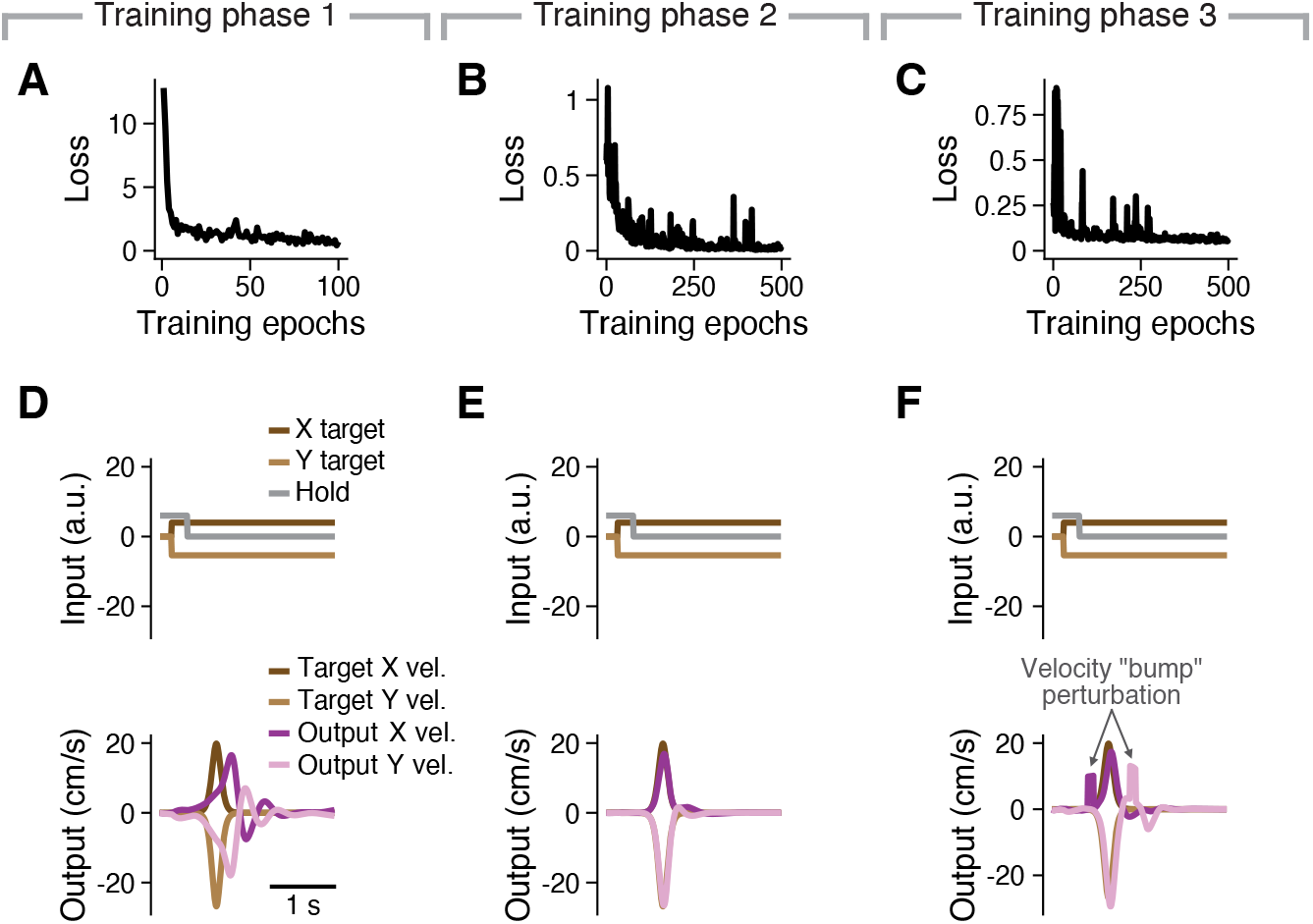
Initial training protocol. The initial training of the RNN is divided into three phases. In the first phase, we kept the feedback weights *W* ^*fb*^ fixed and at small initial values (A,D). In the second phase, we lifted that constraint, and all model parameters became plastic (B,E). In the third phase, we introduced random velocity perturbations in 75 % of trials (C,F). **A-C**. Training loss for each of the three training phases. **D-F**. Input (top) and output (bottom) for an example test trial for each of the three training phases.

**Figure S2:**
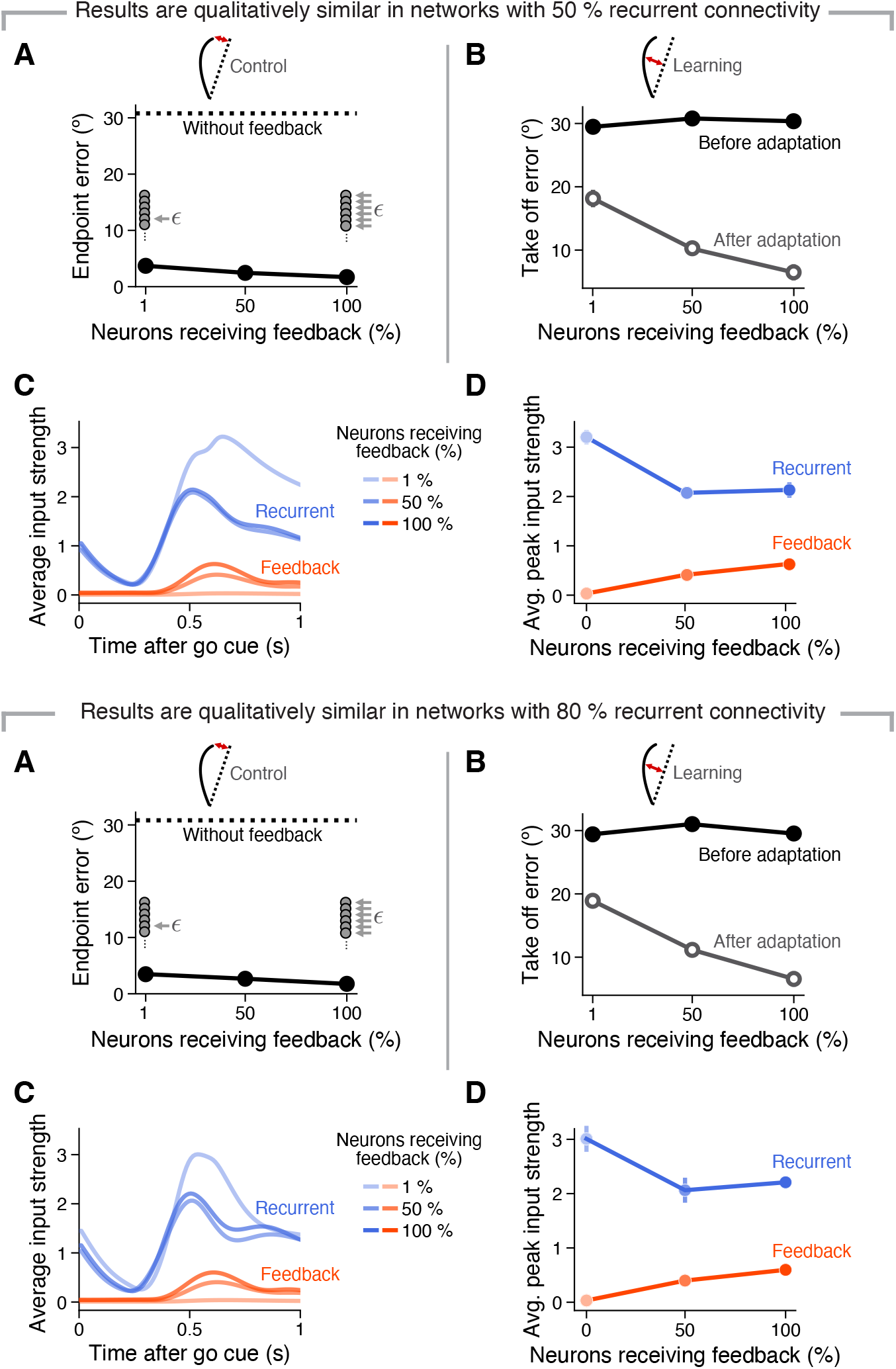
Effective feedback-based motor control and adaptation can both be achieved with sparse feedback for varied degrees of recurrent connectivity. Simulation results for networks with 50% (**A-D**) or 80% (**E-H**) recurrent connection probability. **A**. Angular error between produced and target position at the end of the reach immediately after onset of visuomotor rotation onset for networks with different percentages of neurons receiving afferent feedback (black markers), including no feedback (dashed line). Lines and error bars, mean and s.d. across ten networks. **B**. Take-off error at visuomotor rotation onset (solid circles), and after adaptation (empty circles) for networks with different percentages of neurons receiving afferent feedback. Lines and error bars, mean and s.d. across ten networks. **C**. Average recurrent (blue) and feedback (red) inputs to an RNN neuron before adaptation. Average input strength is defined as the mean across incoming signals and neurons. Legend, percentage of neurons receiving feedback. **D**. Average magnitude of the peak input strength of the recurrent (blue) and feedback (red) inputs before adaptation. Same colour scheme as in C. Lines and error bars, mean and s.d. across ten networks. Data in E,F,G,H are presented as in A,B,C,D.

